# Optimized guide RNA selection recommendations for using *sp*Cas9 gene editing in human hematopoietic stem and progenitor cells

**DOI:** 10.1101/2020.12.18.423329

**Authors:** Han J.M.P. Verhagen, Carlijn Kuijk, Anne M. Kokke, Laurens Rutgers, Santhe A. van der Meulen, Gerard van Mierlo, Carlijn Voermans, Emile van den Akker

## Abstract

Ribonucleoproteins (RNPs) are frequently applied for therapeutic gene editing as well as fundamental research, because the method is fast, viral free, and does not rely on clonal selection.

We evaluated various parameters to genetically engineer human hematopoietic stem progenitor cells (HSPCs) using *sp*Cas9-RNPs and achieve gene editing efficiencies up to 80%. We find that single guide RNA (sgRNA) design is critical to achieve high gene editing efficiencies. However, finding effective sgRNAs for HSPCs can be challenging, while the contribution of numerous *in silico* models is unclear.

Here we established a time- and cost-efficient *in vitro* transcribed sgRNA screening model in K562 cells to identify sgRNAs that are effective in HSPCs using RNP delivery. We show that this simple screening method outperforms all *in silico* prediction models. Our data demonstrates that most *in silico* sgRNA prediction models are ineffective and we make recommendations to potentially improve their accuracy.

We report that gene editing is equally efficient in distinct CD34^+^ HSPC subpopulations. Furthermore, no effects on cell proliferation, differentiation or *in vitro* hematopoietic lineage commitment were observed. Finally, no upregulation of p21 expression was found, suggesting unperturbed HSPC homeostasis.

**Key points:**

- *In vitro transcribed* single sgRNAs (IVTsgRNA) screening in K562 outperforms *in silico* modeling
- Hematopoietic stem and progenitor cells are equally targeted by Ribonucleoproteins (RNPs)
- Hematopoietic stem and progenitor cells show no induction of p21 expression or effects on differentiation, proliferation and lineage commitment

**Graphical abstract:** 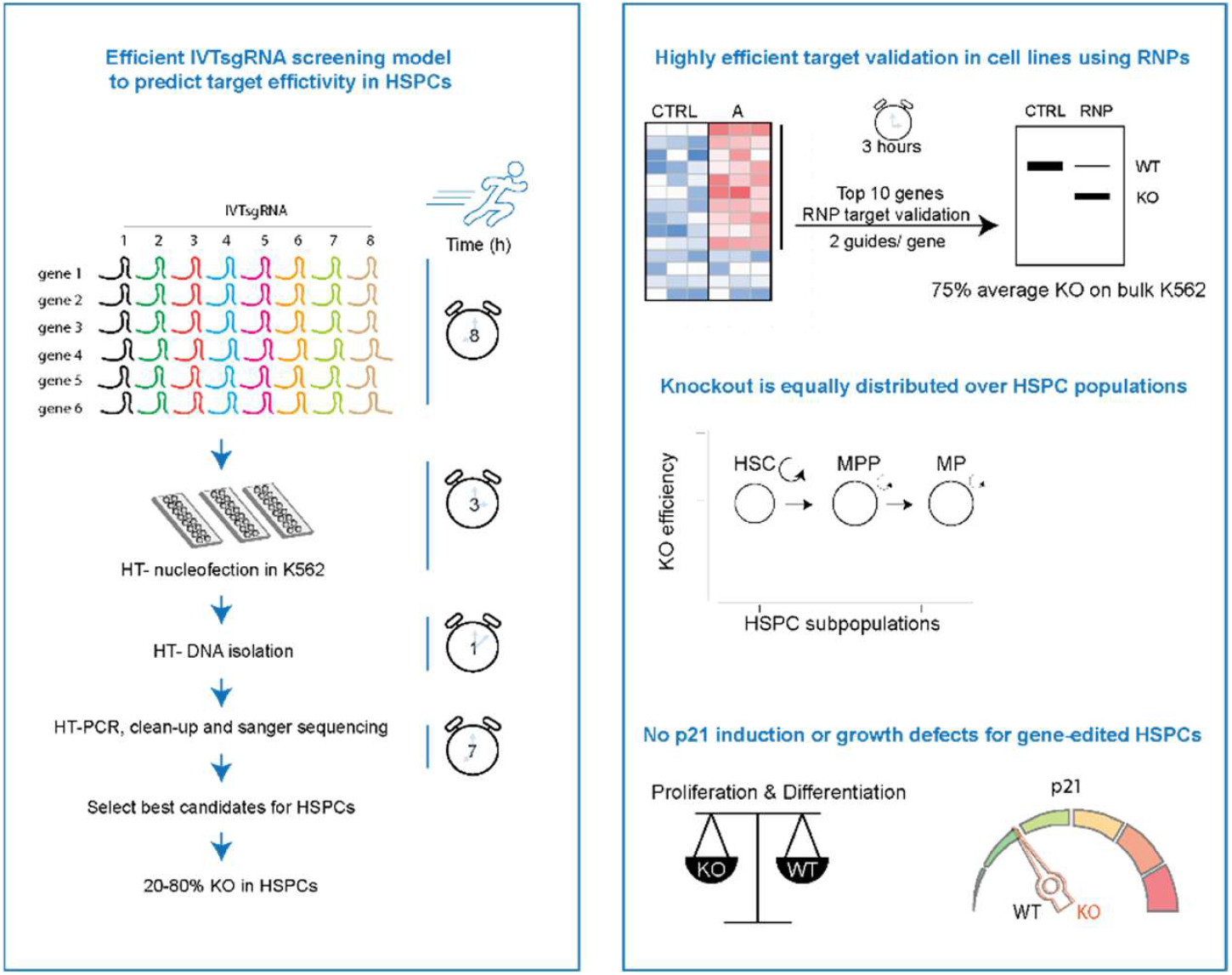

## Introduction

Genome editing is a powerful tool to study gene function and correlation, and the fast turnover time of generating knockout cells makes this technique already indispensable for most research fields. Various genome editing technologies have been developed in the past decade, including zinc-finger nucleases (ZFNs) (1,2), transcription activator–like effector nucleases (TALENs) (3,4) and the RNA-guided Clustered Regularly Interspaced Short Palindromic Repeats (CRISPR)/CRISPR-associated (Cas) systems (5,6), A variety of Cas proteins from multiple species have been identified (7), with *Streptococcus pyogenes* SpCas9 (*spCas9*) and modifications thereof being the most developed gene-editor to date with several updates that provide distinct features (8-12).

In mammalian cells, *spCas9* and single guideRNA (sgRNA) were initially co-expressed using transfected plasmid DNA in immortalized cells followed by clonal selection (13), Despite its success in immortalized cells, clonal selection hampers the applicability to use gene editing in primary human HSPCs as the expansion potential on single clones is limited both *in vitro* and *in vivo*. Highly efficient gene editing on bulk HSPCs can potentially overcome these limitations and can make gene editing an indispensable tool for fundamental, translational and therapeutic purposes. Nonetheless, achieving high genome editing efficiencies in primary hematopoietic cells remains challenging (14).

Previously, several optimization steps were made e.g. direct *spCas9* protein delivery, which outperforms *spCas9* delivered as mRNA or expressed through plasmid DNA (15). In addition, various *spCas9* modifications revolve around optimizing nuclear localization signals (NLS), which can improve the delivery of RNPs to the nucleus (16,17). RNP delivery can be performed using iTOP (18) or more commonly reported by nucleofection (14-19). For nucleofection, various buffers have been reported that potentially could improve efficacy or reduce the costs for RNP based delivery on a large scale (14-20).

For RNP based gene editing, sgRNAs were initially *in vitro transcribed* (IVT). However, chemical synthesized sgRNA lacking the 5’triphosphate cap (15) outperform IVTsgRNAs by preventing RIG-1 mediated intracellular immune responses in primary cells thereby leading to fewer cellular off-target effects (21,22). Moreover, chemically synthesized RNAs contain modifications that improve RNA stability and have been shown to be more effective compared to IVTsgRNAs (15).

Despite these improvements, sgRNA efficacy must be determined empirically in HSPCs, as it is recommended to test 5-8 sgRNAs for each target (14), suggesting that most sgRNAs are ineffective. Importantly, numerous *in silico* based algorithms were developed to predict on-target sgRNA efficacy (23-30), and can be used to aid in sgRNA selection to genetically modify hematopoietic cells. Nonetheless, it is unclear how these algorithms translate to RNP based delivery and which algorithm is most accurate to predict activity in HSPCs.

CRISPR-*spCas9* induced double strand breaks in HSPCs can potentially lead to impaired proliferation and differentiation as previously shown for retinal epithelial cells and human pluripotent stem cells and HSPCs (31-33). Importantly, DNA damage in hematopoietic stem cells, coupled to failure or faulty of repair, can lead to pathology, premature senescence, bone marrow failure and myeloproliferative neoplasms (34,35).

Here we evaluated various parameters that need to be considered for genomic editing in hematopoietic cell lines and human HSPCs including *sp*Cas9 source, nucleofection conditions and sgRNA design. To determine the most optimal sgRNA criteria to genetically modify HSPCs, we have setup a fast IVTsgRNA screening protocol in the K562 cell line that more closely relates to the hematopoietic system compared to previously reported algorithms (23-30). By testing >90 different sgRNAs using RNP-based delivery we demonstrate that IVTsgRNA screening in K562 is effective to predict RNP based genome editing in human HSPCs and outperforms *in silico* sgRNAs selection tools that are currently available. From the 10 *in silico* algorithms that are available we find that the Wang-score (36) is most accurate to predict effective sgRNAs to genome edit HSPCs but inferior to the cellular screening method shown here. RNP-induced gene editing did not lead to p21 upregulation nor HSPC proliferation or differentiation defects.

## Results

### Efficient gene editing in K562 is dependent on *spCas9* NLS variation and nucleofection buffer

To setup a model system for IVTsgRNA screening we started by optimizing gene editing in the hematopoietic cell line K562. *spCas9* purification was performed (**Supplemental figure S1A-F**), and the commercially available cell line kit, suited for K562 cells, was compared to the previously reported homemade K562 cell line solution (14). K562 cells were nucleofected with an ATTO-labelled ribonucleoprotein (RNP) complex targeting CD33. Nucleofection using this homemade solution resulted in significant higher transfection of RNPs (**Figure 1A**) and increased knockout efficiencies with no significant differences in cell survival (**Figure 1B**), compared to the commonly used Amaxa kit. Overall, this leads to significant different nucleofection scores that takes survival and knockout into account (**Figure 1B, right panel**). Importantly, the homemade buffer is low cost and thereby ideal for high-throughput applications or bulk experiments, and can be stored for at least 12 weeks without loss of activity (**Supplemental Figure 1G**).

**Figure 1.**
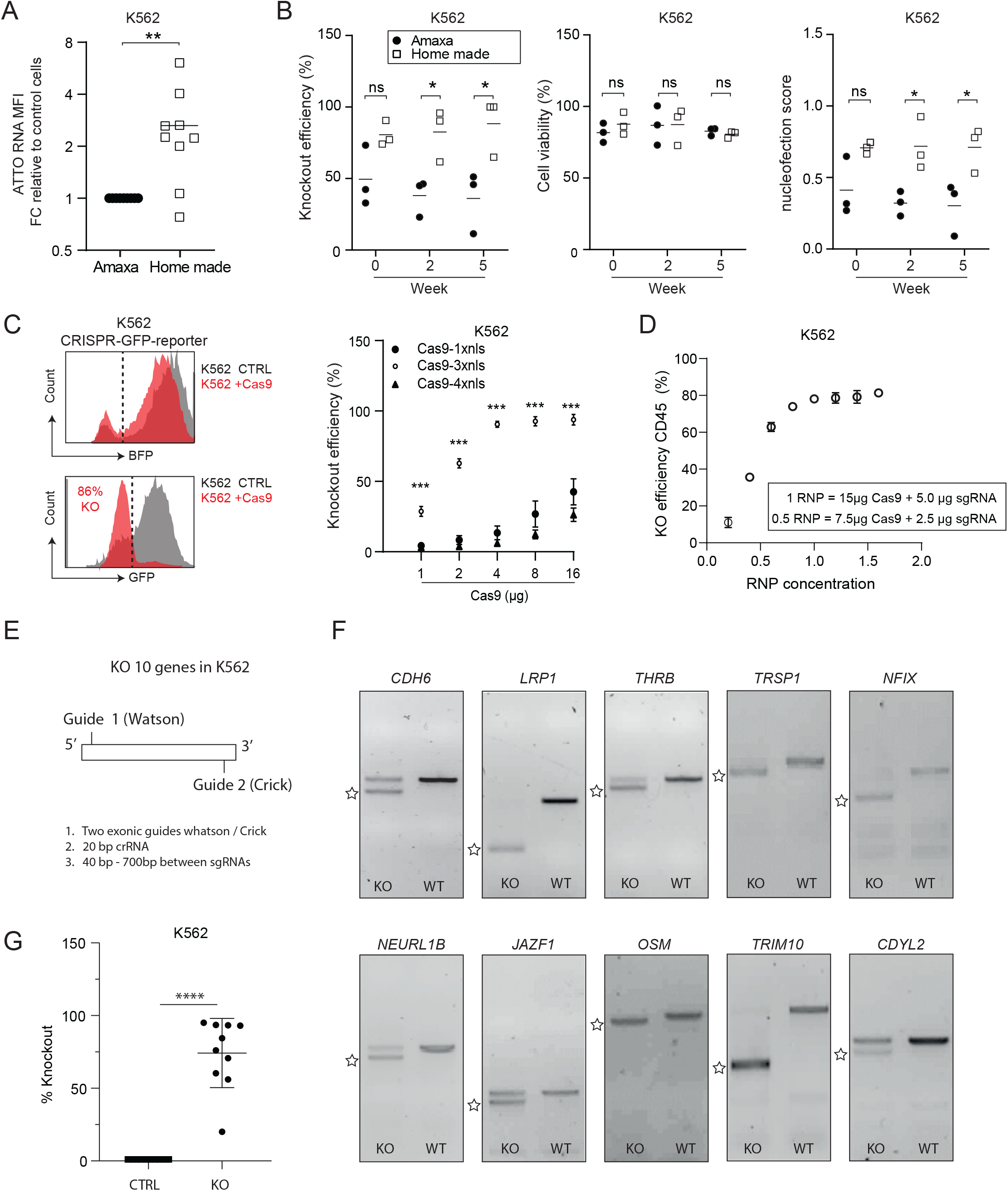
Efficient gene editing in K562 is dependent on *spCas9* NLS variation and nucleofection buffer. **A**) 1-RNP unit of SpCas9-1xNLS-sgCD33-ATTO was nucleofected using three independent Amaxa kits and compared to the homemade K562 nucleofection buffer. Each point shows the relative amount of labeled RNA molecules detected at Day-1 using flow cytometry and samples were compared with an unpaired Student’s t-test. **B**) As Amaxa kits expire within 3 months after opening, we evaluated the performance over time as indicated by week numbers. Cell survival and knockout efficiency were measured after 1 and 4 days respectively post nucleofection using flow cytometry. Each point represents the average of a duplicate and samples were compared using a paired two tailed t-test (**C**) K562-CRISPR-GFP-reporter cells were nucleofected with *spCas9* and analyzed for GFP and BFP expression after 4 days. Data points show the mean ±SD of a n=3 and comparison was made using a One-way ANOVA test followed by Tukey’s post-hoc test. (**D**) RNPs targeting CD45 were tested in various concentrations and KO was measured at day 4 after nucleofection using flow. Data points show the average ±SD of a n=2. (**E**) 20 SgRNAs (2-guide per gene) were designed (supplementary table 1.0) for target validation in K562 cells using Broad institute sgRNA design. (**F**) K562 cells were nucleofected with 1xRNP per sgRNA and DNA was isolated 3-days post nucleofection. Images show PCR products where the KO band is indicated with an asterisk. (**G**) Quantification of KO band/WT product by Image J software and comparison was made using a Student’s t-test (*<p0.05,**p<0.01, ***p<0.001, ****p<0.0001). sgRNA sequences and primers are in supplementary table 1.0.

Next, we evaluated recombinant *spCas9* proteins with different nuclear localization signals (NLS). NLS number and sequence can substantially influence genome editing efficiencies (16,17). Therefore, K562 cells were lentivirally transduced to stable co-express a bicistronic BFP-GFP-reporter and sgRNA against GFP (**Figure 1C**). All transfected *spCas9* proteins generated knockouts in a dose dependent manner. However, 3xNLS-*spCas9* (17) is significantly more potent compared to 1xNLS (43) or 4xNLS (16). This indicates that fusion of specific nuclear localization motifs can significantly enhance genome editing efficiencies. Of note, KO efficiencies for 3xNLS-*spCas9* reached a plateau in K562 between 0.6 and 1.0 RNP units (**Figure 1D**). We next determined if this in-house generated low-cost RNP nucleofection protocol is suitable for high-throughput target validation. For 10 genes, two sgRNAs on opposite strands were designed (**Figure 1E, supplementary table 1.0**), which has previously been shown to enhance genome editing efficiency (44) Strikingly, we find that 9/10 conditions show significant gene alterations, with efficiencies ranging between 50-100% (**Figure 1F-G**), providing evidence that this RNP delivery system is highly suitable for fast and efficient target validation of multiple genes.

### Highly efficient genome editing in human HSPCs is mainly dependent on sgRNAs design

Encouraged by the results in K562, we subsequently aimed to evaluate the translation for genome editing in human HSPCs, and therefore assessed various nucleofection conditions. Several programs are currently reported to transfer RNPs into HSPCs (14-17,39), however, it is unclear which program works best. Here we confirmed that nucleofection program EO100 works significantly better compared to other previously reported programs, indicated by the nucleofection score that takes both survival as well as transfection efficiency into account (**Figure 2A**). Next, it was determined which nucleofection buffer is superior to transfer RNPs into HSPCs, using a previously validated sgRNA targeting CD45 (39) in combination with *sp*Cas9-3xNLS. Of note, we validated that *spCas9*-3xNLS outperforms *spCas9*-1xNLS in HSPCs (data not shown). This revealed that Amaxa P3 provides significant higher gene editing efficiencies compared previously reported buffers (14-20) (**Supplementary figure 2A**), however, differences were generally small and homemade nucleofection buffers (14-20) might alternatively offer an advantage to maintain cost-efficiency. Then it was evaluated which RNP concentration is most optimal to gene edit human HSPCs using CD45 targeting RNPs on distinct HSPC donors. This experiment revealed that 2-3 RNP units is most efficient to achieve high gene editing activities in HSPCs (**Figure 2B**), while cell viability and immunophenotypes of different HSPCs subsets remained similar (**Figure 2C**). Surprisingly, no effect on cell survival or gene editing activity was observed when different cell numbers were target with a fixed amount of RNP molecules (**Figure 2D**). Together these data suggested that high gene editing efficiencies can be achieved in HSPCs. To test if we can reach similar gene editing activities for other sgRNAs, we selected four sgRNAs to target *RUNX1* and tested these sgRNAs in pairs on HSPCs (**Figure 2E**). Here we observed that gene editing efficiencies were generally low and agree with previous findings that most sgRNAs are ineffective in HSPCs using RNP delivery (14).

**Figure 2.**
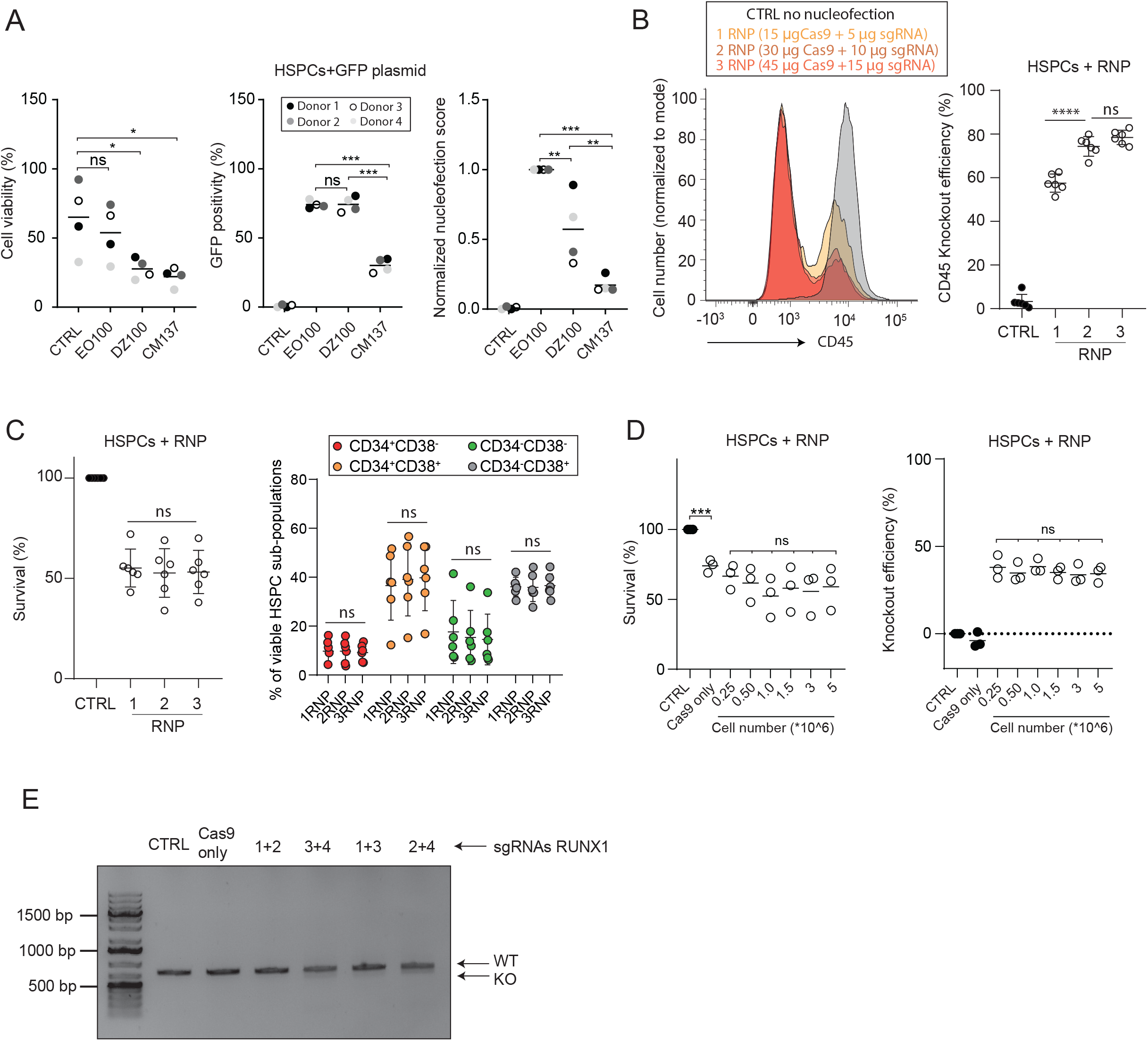
Highly efficient genome editing in human HSPCs is mainly dependent on sgRNAs design. **A)** hCD34^+^ HSPCs were nucleofected with Amaxa P3 buffer and 1 µg of pMax-GFP plasmid on indicated programs. Cell survival and GFP-positivity were measured using flow cytometry 1-day post electroporation. Nucleofection score = % viable cells * % GFP^+^ cells normalized to the best performing program. **B**,**C**) hCD34^+^ HSPCs were nucleofected with distinct concentrations (specified in figure) of spCas9-3xnls molecules targeting CD45. Cell survival was measured on day 1 post nucleofection and KO as well immunophenotype at day 4. **D**) Indicated cell numbers were nucleofected with 1xRNP targeting CD45 or SpCas9 only. Survival was measured 24 hours post nucleofection and KO after 4-days. Statistics were calculated using a One-way ANOVA with Tukey’s post hoc test. (*<p0.05, **p<0.01, ***p<0.001, ****p<0.0001). **E)** Agarose gel showing *RUNX1* PCR products derived from DNA isolates of gene edited HSPCs. Numbers indicated the two sgRNAs that were used per condition and sequences are in supplementary table 1.0.

### IVTsgRNA K562 screening system accurately predicts sgRNA activity for HSPCs

Because most sgRNAs are ineffective in HSPCs using RNPs, effective sgRNAs need to be identified by either screening, or *in silico* modeling. Screening with synthetic sgRNAs is expensive and affects the stocks of precious CD34^+^ HSPC resources. While the contribution of numerous *in silico* models is unclear, as none of these algorithms is validated for RNP-based gene editing in HSPCs.

To overcome this problem, we developed a cost-efficient and rapid *in vitro* transcribed IVTsgRNA screening assay in hematopoietic K562 cells. We hypothesized that this model is more relevant to identify effective sgRNA then *in silico* modeling, as *in silico* models are mainly based on lentivirally expresses sgRNAs in epithelial cells (summarized in supplementary table 1.1). IVTsgRNAs can be screened in bulk as they are low in costs and is a potential match with K562 cells that previously showed no response to IVTsgRNAs (21).

For 6 genes, 5-10 sgRNAs were designed (supplementary table 1.2) and PCR templates including a T7 promoter were generated and used to express IVTsgRNAs (**Supplemental Figure S3A-C**). Screening with this panel of IVTsgRNAs in K562 revealed for most genes at least one candidate that potentially predicts effectivity in HPSCs (**Figure 3A)**. To test our hypothesis, the efficacy of the top performing sgRNA candidates was then monitored in primary HSPCs (**Figure 3B**). This revealed an average indel score of 37%, and we concluded that this IVT-screening system in K562 can be used to identify effective sgRNAs to gene edit HSPCs. Note that synthetic sgRNAs were used for HSPCs as IVTsgRNAs upregulated the RIG-one pathway and induced differentiation in primary hematopoietic cells.

**Figure 3.**
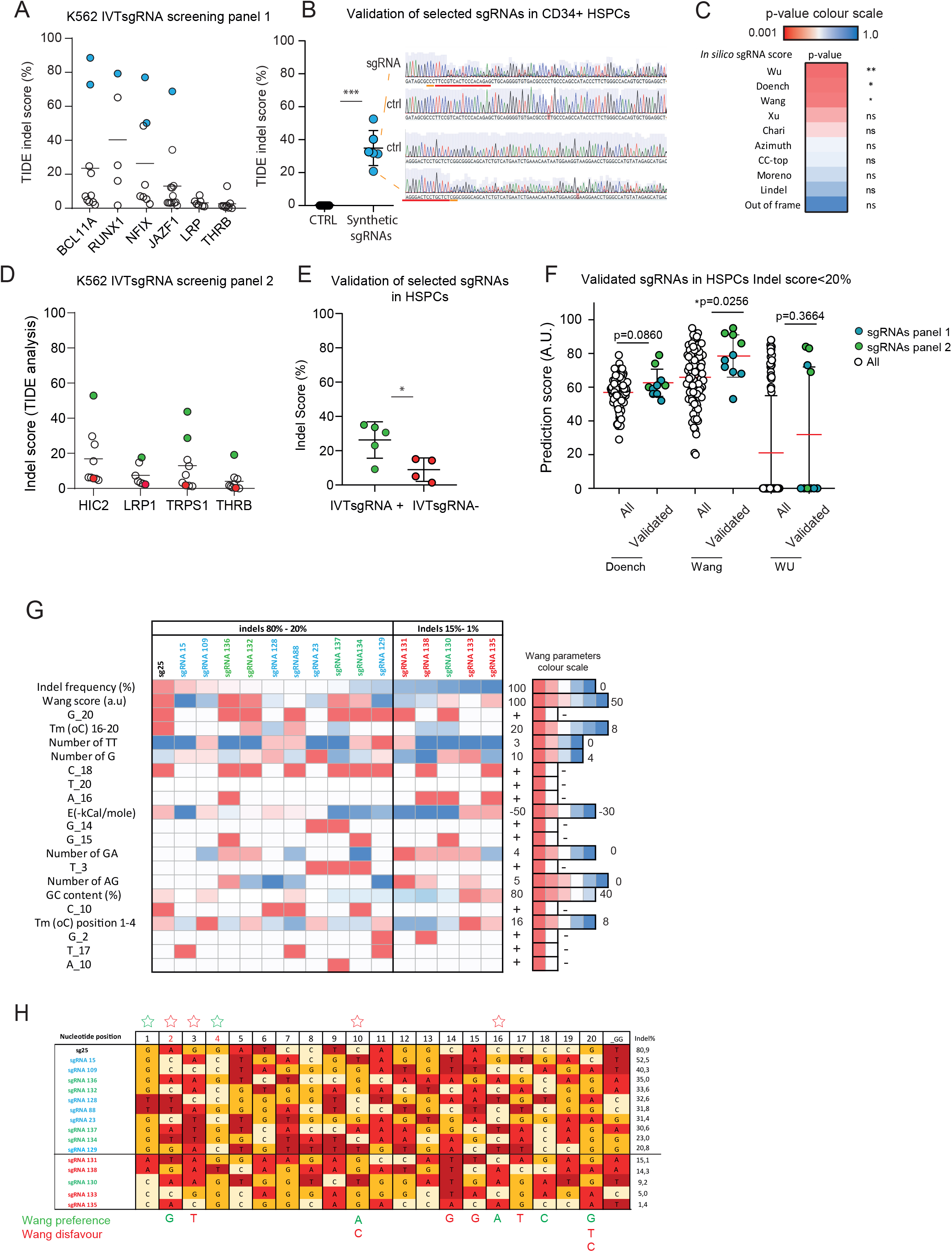
IVTsgRNA K562 screening system accurately predicts sgRNA activity for HSPCs. **A**) 58 IVTsgRNAs were generated and tested in K562. Tide Indel scores were calculated on day-3 post nucleofection. Numbers in graph indicate unique institute codes and sgRNA sequences can be found in supplementary table 1.2. **B**) Six top candidate sgRNAs were ordered synthetically and were tested in CD34^+^ HSPC cells for activity using 2xRNP. Tide indel Scores were determined 3-days after nucleofection. Two example chromatograms for sg23 and sg88, the guide is indicated in red and PAM site in orange. **C**) in HSPC validated sgRNAs were analyzed retrospectively for in silico prediction scores (graphs in supplemental Figure 3). Heatmap represents the summary of the statistic analysis comparing the validated sgRNAs with all the 58 sgRNAs. **D**) Panel 2 shows the IVTsgRNA tested in K562 that were enriched for high Wang, Doench and Wu scores, each dot show a distinct sgRNA. Top performing sgRNAs (green) and poor performing sgRNAs with a similar in silico score (red) were selected for further analysis. **E**) CD34^+^ HSPCs from the same donor as used in panel 1, were targeted with synthetic sgRNAs and delivered as 2xRNPs. * ≥0.05 from a Student’s t-test. **F**) Validated sgRNAs from panel 1 (blue) and panel 2 (green) were analyzed for the top 3 performing *in silico* models and comparison was made using a two-tailed t-test except for the WU score where an unpaired Mann Withney test was used because the abnormal distribution. **G**) All parameters from the Wang score were analyzed for the tested sgRNAs and illustrated in a heatmap. We defined ineffective and effective sgRNAs with Indel score <15% and >20% respectively **H**). Detailed sequence analysis of sgRNAs that we have tested in HSPCs and comparison to Wang algorithm.

With this panel of validated sgRNAs, we addressed the question which *in silico* algorithm would retrospectively have predicted most accurately these validated sgRNAs (**supplementary Figure 3D)**. Here we identified Wang, Wu and Doench’16 score as most predictive to enrich for sgRNAs that are effective in HSPCS (**Figure 3C**).

Based on these observations a second panel of IVTsgRNAs was designed, this time IVTsgRNAs were selected with high Doench-Wang and Wu scores (supplementary table 1.3). Screening in K562 revealed several potential effective sgRNAs (**Figure 3D**). Given that most of the sgRNAs are ineffective we hypothesized that either *in-silico* prediction scores are ineffective, or that the IVTsgRNA screening model results in false negatives. For this reason, top candidates (green) were validated in HSPCs (**Figure 3E**), however, this time also sgRNAs were tested in HSPCs with a similar Doench, Wang and Wu score that were found ineffective in the K562 IVTsgRNA screening model (red). The sgRNAs identified in the K562 screening showed a significant higher Indel score in HSPCs compared to their sgRNAs that were assigned as ineffective. This proves that screening of IVTsgRNAs in K562 is a predictive model for using synthetic sgRNAs in HSPCs. To find which algorithm would have been retrospectively most effective, all validated sgRNAs with an indel score higher than 20% were plotted and compared to the complete data set (**Figure 3F**). This analysis revealed that the Wang score is most predictive, however, given that effective sgRNAs can be found within the complete range distribution of the Wang score we conclude that cellular screening is still superior. No correlation was found between the Wang score and indel formation (**supplemental Figure S3E**). In search for an explanation why solely *in-silico* modeling is not sufficient, all parameters from the Wang score were analyzed for the validated sgRNAs and compared to their counterparts that were ineffective (**Figure 3G**). Moreover, the exact sequences of the validated sgRNAs were aligned (**Figure 3H**) and compared to the preferred and disfavored nucleotides that are used in the Wang algorithm. This analysis explains partly why *in silico* prediction is ineffective. Note that a majority of the effective guide RNAs do not align with the criteria of Wang (indicated by red asterisk) and also trends were observed in our data set that were not previously reported or included in the Wang score (Green Asterisk).

### Genome editing efficiencies are equally represented within distinct human HSPC subpopulations

HSPCs can be divided in different subpopulations with distinct self-renewal and or differentiation properties (45). Knockout efficiencies in CD34^+^CD38^-^ immature hematopoietic stem cells, CD34^+^CD38^+^ hematopoietic stem and progenitor cells and CD34^-^CD38^+^ myeloid lineage committed cells were similar (**Figure 4A,B**). Further subdivision of the CD34^+^CD38^-^ cells using CD45RA and CD90 into immature hematopoietic stem cells (46,47) demonstrated that genome editing efficiency was similar to downstream progenitors, albeit that events assayed was rather low (Between 150 and 4200, **Supplemental Figure 4**). Overall, we show that genome editing of HSPCs is consistent between donors and that genome editing efficiencies are similar between CD34^+^ HSPC subpopulations.

**Figure 4.**
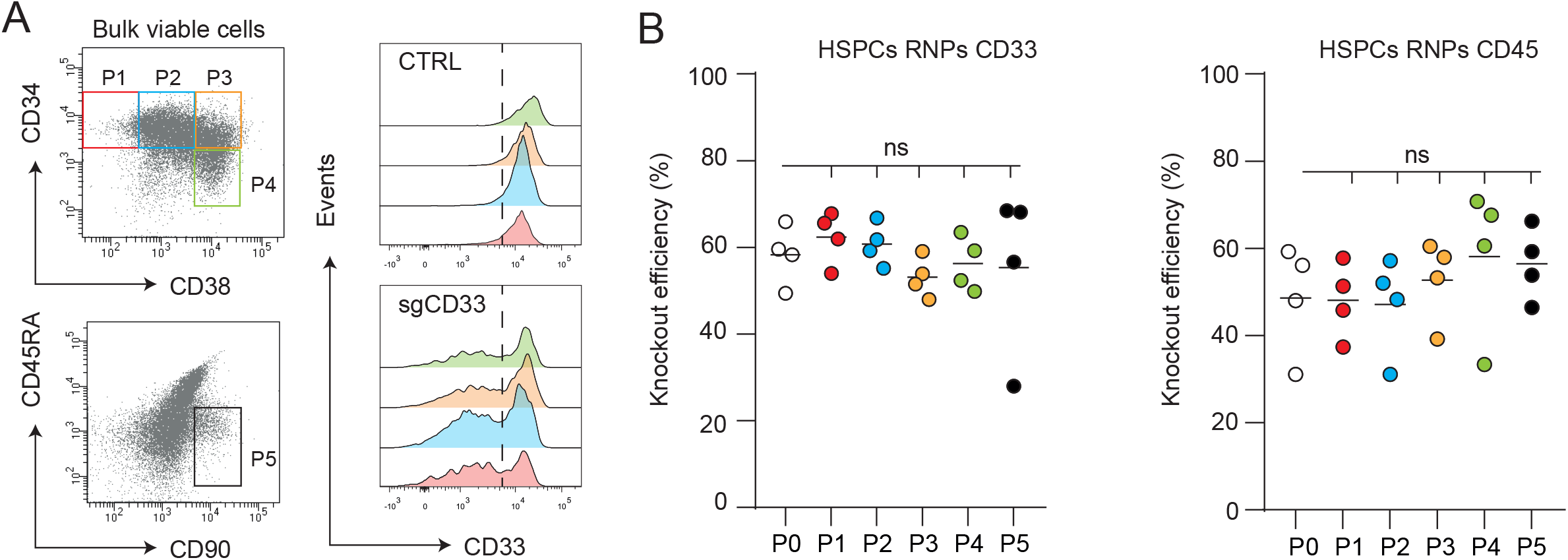
Genome editing efficiencies are equally represented within distinct human CD34^+^ HSPC subpopulations. **A**) FACS plots showing the distribution of distinct hCD34^+^ HSPCs populations from 4 different donors that were nucleofected with 1xRNP targeting CD33 or CD45. At day 5 the KO was determined on indicated CD34^+^ populations, P1 - P4 were gated on the viable cell fraction, P5 fraction was analyzed on the P1 fraction. **B**) The KO from the 4 different donors was calculated for CD33 and CD45. Statistics were calculated using a One-way ANOVA with Tukey’s post hoc test. (*<p0.05, **p<0.01, ***p<0.001, ****p<0.0001).

### p21 is not induced after highly efficient genome editing in human HSPCs

CRISPR-*spCas9* induces double strand breaks, which has been reported to lead to p53 DNA damage response pathway activation and transcriptional activation of Cyclin dependent kinase inhibitor p21CIP1/Waf1 regulating G1 cell cycle progression. Genome editing may potentially lead to enrichment of cells that are, to a variable degree, defective in their p53 response (31-33). For this reason, we sought to determine if human HSPCs upregulated p21 when targeted with spCas9 RNPs. Incubation with Nutlin, an inhibitor of the interaction between MDM2 and p53 thus stabilizing p53, lead to a stark decrease in cell survival in wild type p53 (p53^wt^) AML cell lines but not in p53 mutant cell lines (**Figure 5A**). Nutlin induced p21 expression specifically in p53^wt^ cells but not in p53 mutant cells indicated by Western-blot (**Figure 5B)**. Note that ML2 is weakly positive, indicated by an increased intensity image **(supplemental S5A**). Induction of p21 was confirmed by flow cytometry and expression levels correlated with protein levels (**Figure 5C**). Next, we assayed if CRISPR-induced DSBs in p53^wt^ cells lead to reduced cell survival as observed upon stabilizing p53 by Nutlin. The bicistronic BFP-GFP-sgRNA CRISPR reporter gene was stably expressed in the 5 different cell lines (three p53^wt^ and three p53 mutant lines; **supplemental Figure S5B**). Of note, nucleofection programs were optimized for all indicated cells (**Supplemental Figure S5C-E**). The BFP-GFP-CRISPR reporter lines were co-cultured with their wildtype counterparts (1:1 ratio) to perform competition experiments. We hypothesized that wild type cells outcompete the BFP^+^ cells if DSBs induce p53/p21-mediated cellular senescence (**Figure 5D**). *spCas9* transfection led to a significant GFP reduction in all cell lines, however, no correlation between GFP knockout and the loss of BFP^+^ cells was observed (**Figure 5E**). This suggested absence of p21-induced senescence. Next, the responses to RNP-induced DSBs in purified mobilized peripheral blood CD34^+^ HSPCs were measured. As expected, HSPCs upregulated p21 (**Figure 5F**) upon incubation with the p53 stabilizing agents Nutlin, resulting in significant cell death after 16 hours (**Supplemental Figure 5F**). This indicates that HSPCs are responsive to p53 activation and react by upregulating p21. Next, we induced multiple DSB breaks using different RNP complexes. Knockout efficiency assayed after 4 days was 60% for CD33 and 50% for CD45 (**Supplemental Figure 5G**,**H**). The double knockouts resulted in a somewhat lower knockout efficiency of approximately 20%, (**Supplemental Figure 5I**). In contrast to treatment with the p53 stabilizing agent Nutlin, both the single and double RNP complex nucleofected CD34^+^ cells did not show significant upregulation of p21 (**Figure 5G**). This suggests that p53 is either not activated due to RNP-induced DSBs or that this activation remains too low to detect using p21 as a readout.

**Figure 5.**
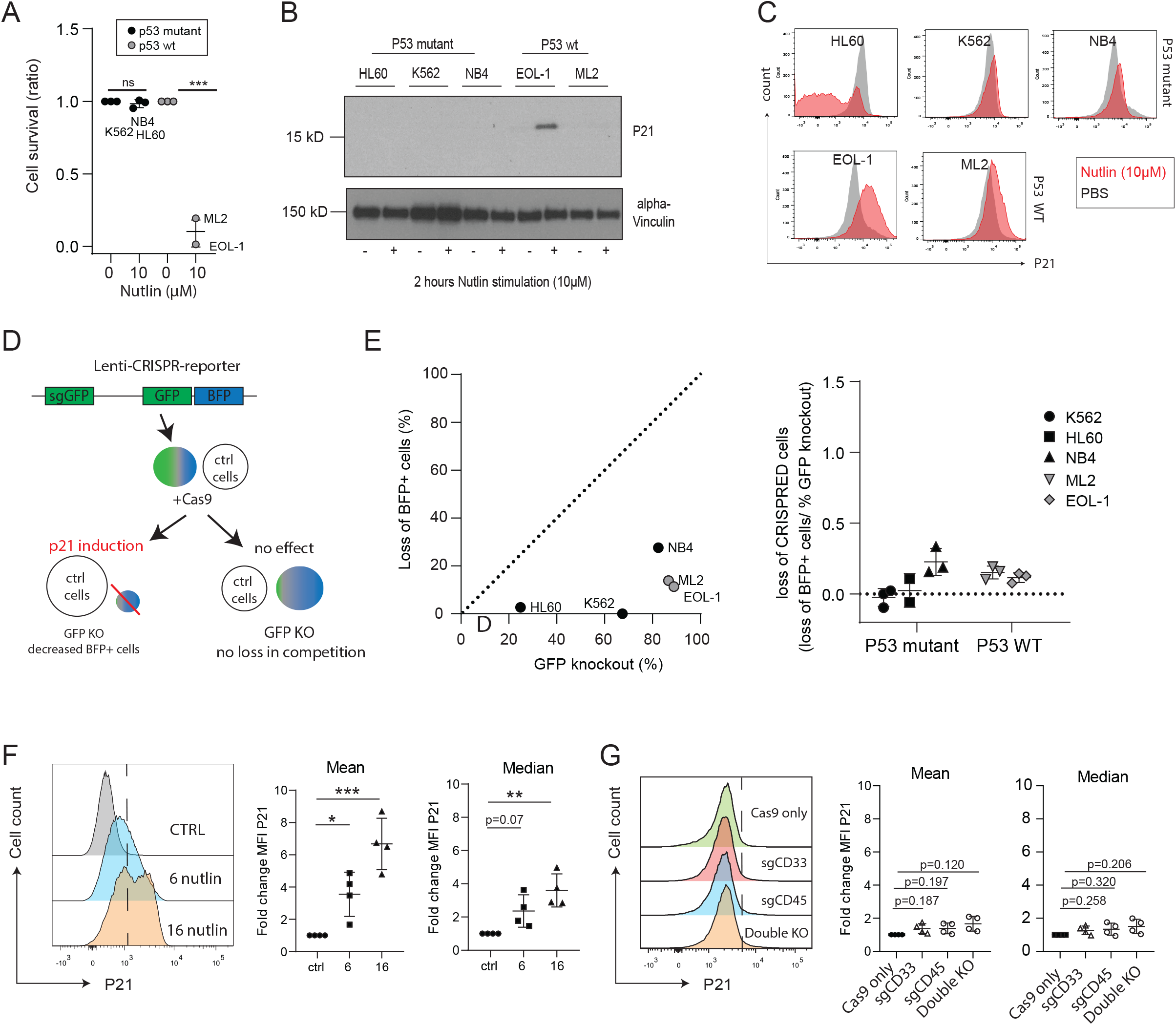
p21 is not induced after highly efficient genome editing in human CD34^+^ HSPCs. **A**) AML cells were culture on 0.5×10^6^ cells/ml and incubated with 10 µM Nutlin for 72 - 96 hours. Cell survival was analyzed using flow cytometry and graph presents the data from a n=3. Significance was calculated using a paired Student’s t-test. **B**) 0.5⨯10^6^ cells/ml were incubated with 10 µM Nutlin for 2 hours and lysates were analyzed for p21 activation (note extra exposure of Western Blot in supplementary figure 5A for increased contrast). **C**) Simultaneously to the Western Blot from Figure B, cells were analyzed using flow cytometry for p21 expression on the viable cell population. **D**) Model explaining competition assay designed for leukemic cell lines. 100% GFP^+^ cells were mixed to their WT-counterparts (1:1 ratio) and hence nucleofected with SpCas9. In case p21 induces cellular senescence, BFP+ cells will lose the competition from WT cells over time. **E**) Representative graph from a single GFP competition experiment where cells were analyzed for BFP and GFP levels using flow at day 4-post nucleofection with SpCas9 (left panel). The loss of genetically modified cells (right panel) was calculated by (BFP^+^ cell loss/ GFP knockout) (relative to non-nucleofected cells). Graph shows the quantification of the loss of BFP^+^ cells at day 4 post nucleofection from a n=3. Data were analyzed using a One-way ANOVA followed by Tukey’s post-hoc test. **F**) hCD34^+^ HSPCs were cultured in presence of 10 µM Nutlin or PBS and p21 staining was measured using flow after indicated time points. Graphs show the expression of p21 for 4 different donors at indicated time points. Fold changes were calculated relative to control cells, and P values were calculated using a One-way ANOVA test and Tukey’s post-hoc test. **G**) CD34^+^ cells were nucleofected with 1xRNP for CD33, 1xRNP for CD45 and the combination represented as double KO. p21 stainings were analyzed on day-5 after CRISPR. Graphs show the fold change relative to control cells and was analyzed using a One-way ANOVA with Tukey’s post hoc test. (*<p0.05, **p<0.01, ***p<0.001, ****p<0.0001)

### Gene edited HSPCs show no defects on differentiation, proliferation or lineage commitment

*sp*Cas9 potentially hampers cell proliferation or differentiation of HSPCs as suggested previously (33). To test this hypothesis, knockouts were generated in HSPCs and differentiated towards erythroid cell cultures (**Figure 6A)**. As CD45 and CD33 are not expressed on erythroid cells, the knockout dynamics were assayed using an RNP-complex targeting CD44. Using 1-RNP, we observe 40% KO and limited cell death at day 5, indicating fast recovery (**Figure 6B**). Indeed, CD44 knockout was unchanged during erythroid proliferation and cells expanded similarly to control cells (**Figure 6C**). Furthermore, no differences were observed during differentiation indicated by erythroid immunophenotypic staining’s of CD235 and CD71 (**Figure 6D**). The nucleofection process and subsequent RNP-induced DSBs leading to specific knockout of CD33, CD45 or both genes may affect the specific outgrowth of CD34^+^ HSPC to lineage effector cells. However, the frequency of CD13^+^ myeloid cells, CD235^+^ erythroid cells and CD41^+^ megakaryoid/HSPCs did not change between cells that were successfully genome edited or that were nucleofected with *spCas9* only (**Figure 6E**). Importantly, the knockout efficiency within differentiated lineage cells was not altered compared to day 4. This showed that nucleofection as well as DSB induced by RNP complexes do not interfere with the specification to specific myeloid lineages. In addition, the data confirms that knockout of CD33, CD45 or the combination does not intrinsically interfere with differentiation to myeloid lineage.

**Figure 6.**
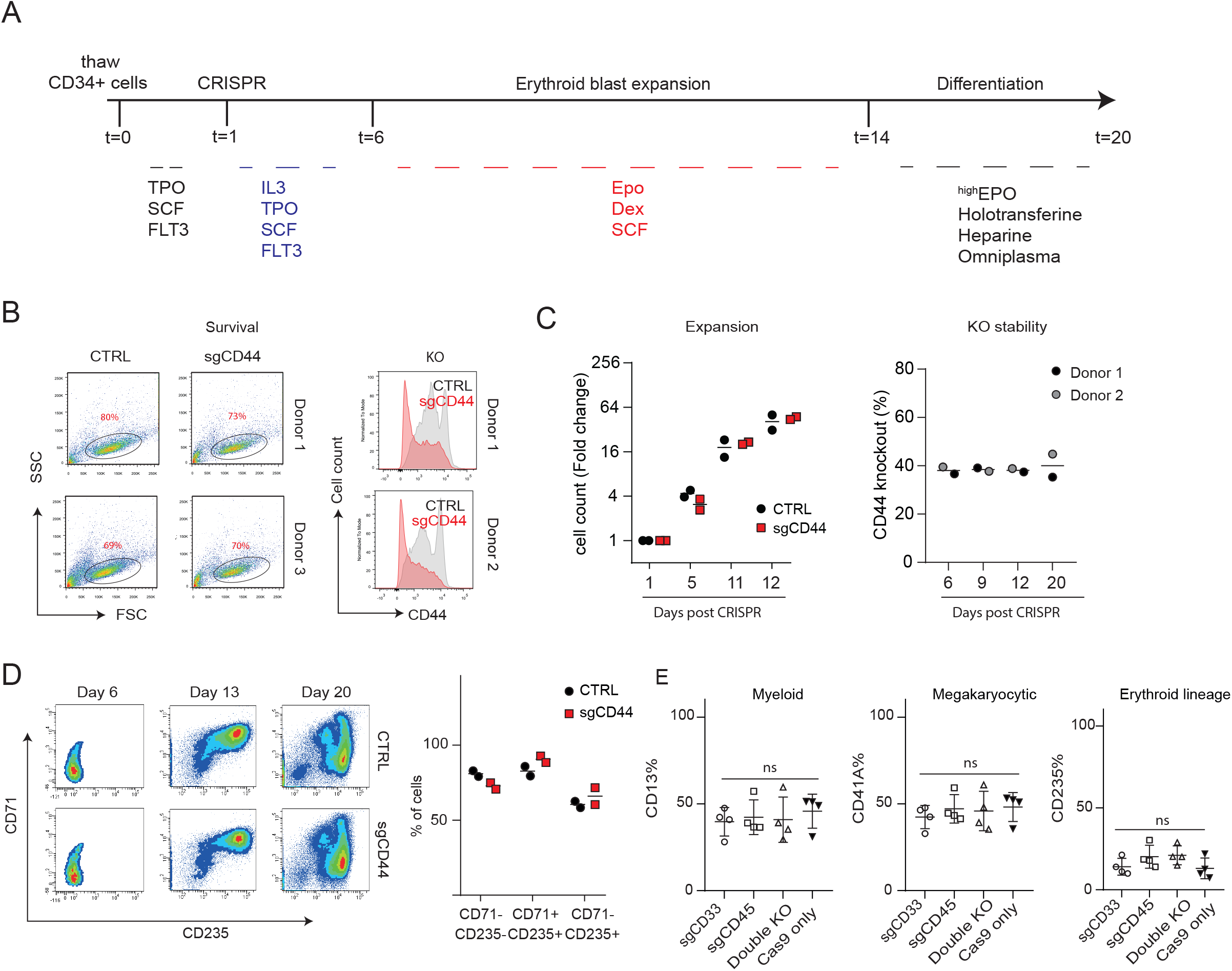
CRISPR-SpCas9 has no effect on the differentiation or proliferation of HSPCs. **A**) Culture scheme that was used to study the growth dynamics of gene edited cells. **B**) Flow cytometric measurement of genetically modified hCD34^+^ HSPCs analyzed for cell survival and KO efficiency on Day-1 and Day-5 respectively. **C**) Expansion was determined using a Casy-counter and fold change was calculated based on cell count at day 1 at the start of the experiment (left panel). KO stability was determined using flow cytometry on viable cells at indicated time points. **D**) Erythroid differentiation was analyzed at indicated time points (example left panel) and quantified for two donors (right panel). **E)** CD34^+^ HSPCs with KO efficiencies of ∼50% (**Supplementary figure 5H**), were analyzed for indicated cell lineage output as determined by flow cytometry, 9 days after CRISPR. On Day 1-4 cells were cultured in media supplemented with a cytokine maintenance cocktail containing SCF (100 ng/ml), FLT3 (100 ng/ml), and TPO (10ng/ml), followed by 5 days cultured in media containing SCF 100 ng/ml, TPO 10 ng/ml, EPO (2IU/ml), IL3 (100 ng/ml), IL6 (100 ng/ml) GM-CSF (100 ng/ml). Data were analyzed using a One-way ANOVA with Tukey’s post hoc test. (*<p0.05, **p<0.01, ***p<0.001, ****p<0.0001).

## Discussion

CRISPR *spCas9* gene editing is a powerful tool to study gene functions in the hematopoietic system. The protocols described here show that an optimized RNP system can result in high gene editing efficiencies in K562, independent of sgRNA selection. Our data shows that the 3xNLS-spCas9 protein was greatly superior compared to 1xNLS (43) and 4xNLS (16) in gene editing within K562 and HSPCs. Concomitantly, 3xNLS-spCas9 was previously shown to outperform 2xNLS*sp*Cas9(17). Surprisingly, similar gene-editing efficiencies for 1xNLS (43) and 4xNLS (16) were observed, where it was previously demonstrated that 4xNLS significantly outperforms 1xNLS and 2xNLS-SpCas9 proteins by local delivery in mouse brain (16). We speculate that the performance of *sp*Cas9 proteins and distinct fused NLS-sites might be cell type dependent.

In general, our results indicate that genome editing in primary human HSPCs is less efficient compared to K562 and other cell lines that we have tested. This might be caused by (i) the nucleofection buffer, which is likely different for K562 compared to primary cells, and (ii) the intrinsic nature of the cells that is clearly distinct for HSPC cells (e.g. DNA repair machinery or nuclear transport of RNPs).

Our data clearly demonstrate that sgRNA selection is of critical importance to generate Indel formations >20% in HSPCs. This study demonstrates that *in silico* prediction models are not predictive while in contrast IVTsgRNA screening in K562 predicts effective sgRNAs for spCas9 in HSPCs. Our data set could disprove several of the suggested nucleotide preferences or disfavors from the Wang score. For instance, position 2 is, according the Wang Score, preferred to be a Guanine while our data sets revealed one sgRNA with a guanine on position 2. We speculate that position 1 is more important like Wang and colleagues showed previously for the high fidelity spCas9 proteins (36). The Wang score furthermore speculates that an Adenine is preferred on position 10 and 16, while this was not observed in our data set. However, it must be noted that the number of sgRNA used in this study may only allow correlations to be drawn.

HSPCs may be more difficult to target due to their active p53 status, as compared to K562. Increased p21 mRNA expression levels were previously observed in the first 24 hours using *sp*Cas9 by making a single DSB in HSPCs on the Y-chromosome of male cells (33). Because the p53 pathway is actively involved in the NHEJ route we hypothesize that HSPCs are possibly more efficient in repairing the DSBs through p53-induced DNA repair machinery and thereby leading to decreased genome editing efficiencies. In fact, this suggests that transient p53 pathway inhibition using GSEA56 (33,48,49) can possibly enhance genome editing activities. It was previously shown that transient p53 inhibition using GSE56 prevents a proliferation delay in genetically modified HSPCs using AAV6 and finally resulted in better engraftment, which was found initially delayed in AAV6 genetically edited cells (33).

In our study, no significant p21 protein upregulation was observed in long term and suggests that gene editing using RNPs in HSPCs is safe. This is further supported by our data as no defects were observed in lineage commitment, proliferation and differentiation. Because gene editing efficiencies are equal over distinct HSPC subpopulations, we speculate that transplantation of cells is feasible and paves the road toward implementation for clinical purposes.

## Materials and methods

All reagents and resources are described in supplementary table 1.0 – 1.5 as well as supplementary materials and methods.

### *sp*Cas9 protein expression and purification

spCas9-3xNLS (Addgene plasmid # 114365) was expressed in Bl21 star cells (Thermo Fisher, # C601003) and purified in a two-step purification method (Histrapp, followed by size exclusion). Protein concentration of spCas9 concentrations were determined in BCA proteins assay, and snap frozen store at -80°C. (see supplementary methods for more detailed protocols and resources).

### IVTsgRNA synthesis

Guide RNAs were designed using CRISPOR (Version 4.98) by selecting guide RNA sequences with the highest specificity score (unless stated otherwise). crRNA sequences were developed as ssDNA oligo’s including a T7 promoter and an overhang sequence resulting in a 56-58 nt oligo (see supplementary table 1.4 for examples, note that 1 or 2 Guanines should be added in some cases). This oligo was annealed and amplified to a fixed 91nt oligo containing the scaffold domain using Taq polymerase (Invitrogen, Cat. #10342046). Transcription was performed using Hi-SCribe T7, NEB E2040 and purification with monarch columns (NEB, T2040). Concentration was measured using Nanodrop. (see supplementary methods for more detailed protocols and resources).

### High-throughput IVTsgRNA screening

3 µg spCas9-3xNLS + 1 µg IVTsgRNA was resuspended in 20µl homemade nucleofection buffer(14) using 0.2×10^^6^ K562 cells. 20 µl of the RNP-K562 cell mixture was nucleofected using FF-120 (Amaxa 4-D) and 80 µl of culture media was added directly after transfection. Without washing, the complete 100 µl nucleofection mix was transferred and cultured in 24-wells plates and cultured in 500 culture media. The complete process of RNP formation, transfection and washing of the strips was performed using a multichannel and takes ∼2-3 hours to transfect 48 sgRNAs. Note that nucleofection strips and cuvettes can be re-used as described previously (14). At least 30.000 cells were used for DNA isolation in HT-vacuum manifold according manufactures conditions (MN 740455.4) that can process up to 96 samples in 1 hour. 2 µl of DNA isolate was used for PCR (primers are in supplementary table 1.4) and PCR cleanup was performed using Exo-SAP (NEB, M0293, M0371). Sanger Sequencing reactions were prepared using (Thermo Fisher # 4337455), and TIDE analysis (37) using standard settings was used to determine the Indel Frequency.

### CD34 isolation and nucleofection

Human material was obtained after informed consent, mobilized peripheral blood (MPB) was obtained from leukapheresis material, and cord blood (CB) was collected according to the guidelines of NetCord FACT (by the Sanquin Cord Blood bank, The Netherlands). Cryopreserved CD34^+^ cells were thawed and cultured in Cellquin media (38) supplemented with a cytokine maintenance cocktail containing SCF (100 ng/ml), FLT3 (100 ng/ml), and TPO (10ng/ml). All cells were pre-cultured for 16-30 hours prior to nucleofection. CD34 cells were collected and resuspended in nucleofection buffer from Amaxa or homemade M1 buffer(14) and nucleofected using program EO100 unless otherwise stated. Directly after nucleofection, room temperature culture media was added and cells were spun to remove residual nucleofection buffer. Cells were cultured with Cellquin media including IL3 (100 ng/ml), IL6 (100 ng/ml), TPO (10ng/ml), SCF (100 ng/ml) at 37°C, 5% CO^2^. Expansion and proliferation was performed as previously described (38).

### RNP formation for synthetic sgRNAs used in HSPCs

Synthetic crRNAs (IDT, Alt-R) and Synthetic scaffold RNA (both from IDT, DNA technologies) were dissolved as 100 µM stocks using ultra-pure RNAse free water. From this stock 2,25 µl crRNA was incubated with 2,25 µl Tracer RNA and incubated at 95°C for 5 minutes. Next the mix was incubated at RT for 20 minutes to allow sgRNA formation. 15µg of spCas9 was added and incubated at RT for 10 minutes to form RNPs. In total this is ∼100 pmol Cas9 + 225 pmol sgRNA, as previously reported (14). All guide RNA sequences are reported in supplementary table 1.0 - 1.4. sgRNA sequences targeting CD33 and CD45 were from (39) and CD44 from (40).

### Cell line culture and nucleofection

K562, EOL-1, NB4 and HL60 were cultured in Roswell Park Memorial Institute (RPMI) medium (130-046-703) containing 10% heat-inactivated FBS, while ML-2 was cultured in RPMI medium containing 15% and 20% FBS respectively. All cell lines, were nucleofected with Homemade nucleofection buffer(14). NB4 and HL60 were nucleofected using program X-01, ML2 and EOL-1 at CA-137 and T-016 for K562.

### GFP competition assays

Cell lines were transduced using RetroNectin (Takara biosciences, Cat T202) according manufacturers conditions. The GFP reporter was a kind gift from Kosuke Yusa (Addgene plasmid # 67980 and BFP^+^ cells were enriched using FACS, in case transduction efficiency was < 90%. Transduced cells were mixed 1:1 with their wild type counterparts prior to nucleofection and the BFP ratio was determined on cells that were not transfected with 5-10 µg Cas9-3xNLS.

### Western Blot and flow cytometry

Lysates for Western-blot were made in a 1% triton buffer including cOmplete protease inhibitor cocktail (Roche 11697498001) and Western-Blot and flow cytometric staining was performed as described previously (41),(42). All antibodies for flow cytometry and Western-Blot can be found in supplementary table 1.5.

## Author contributions

HJMPV, CK, AK, LR, SM, GM designed and performed the experiments. CV provided the CD34^+^ HSPCs. Concepts were formulated by HJMPV, EA and CV. HJMPV, CV and EA wrote the initial manuscript and CK, AK, LR, SM, GM made improvements.

## Acknowledgments

Eelke Brandsma for critically reviewing the manuscript, Niels Geijsen, Patrick Celi, Scott Wolfe, Daniel Bauer and for discussing the protein purification setup.

This work was supported by ZONMW game changer (116004203)

