## Supplemental figures for "Optimized guide RNA selection recommendations for using *sp*Cas9 gene editing in human hematopoietic stem and progenitor cells"

### Supplemental information

### Supplemental Materials and Methods

#### Cas9 Protein purification

pET-NLS-Cas9-6xHis was a gift from David Liu (Addgene plasmid # 62934), pET-21a 3xNLS SpCas9 protein expression was a gift from Scot Wolfe (Addgene plasmid # 114365), 4xNLS-pMJ915v2 was a gift from Jennifer Doudna (Addgene plasmid # 88917). Bacterial expression cultures were performed as previously described[19]. Bacteria were lysed in buffer A (20mM Tris, 0.5M NaCl, 50mM TCEP, 2.5mM MgCl2) + protease inhibitor cocktail (Roche) and 10 mM imidazole. His tag purification was performed using a AKTA purifier (GE, health care) in combination with 5 ml His-purification column (GE Healthcare, 17-5248). Washing and elution was done with buffer A+ 40 mM and 250 mM imidazole respectively. Pooled Cas9 containing fractions were concentrated using Amicon ultra centrifugal 100 kDa cutt-off filters (UFC9100 Amicon) and subsequently loaded on HiLoad 16/600 Superdex 200 PG gel filtration column (GE28-9893-35, GE Healthcare) and purified fractions were concentrated with Amicon UFC9100 in buffer A + 0.5M NSDB-201 (Santa Cruz, sc-202237) and filtered using a Acrodisc Mustang E 0.22 µM Endotoxin removal filter (Pall, MSTG25KIT. Cas9 was snap-frozen and stored in an ultra-low freezer (-80oC). Protein concentration was determined using BCA protein assay including 0.1- 0.5M NSDB-201 to correct for background absorption.

#### IVTsgRNA production and purification

The forward oligo was developed including 1 or 2 extra guanines after the T7 promoter to enhance RNA expression[55], resulting in a 56-58 nucleotide oligo containing the 20nt CRISPR sequence (see also examples Table 1.1). A DNA template was formed by running a 35-cycle two-step PCR (10 seconds denaturation 98°C, 10 seconds elongation 68°C) using standard PCR mix as described below. 5 µl of PCR product was incubated with T7 transcription mix for 16-20 hours according to manufacturer's recommendations (Hi-SCribe T7, NEB E2040). DNAse I treatment using 1 µl of DNAse I (NEB, M0303) was performed for 15 minutes at 37°C followed by 15 minutes of inactivation at 80°C. Next purification was performed by using monarch columns (NEB, T2040). RNA concentrations were estimated using NanoDrop and RNA was diluted to 0.5 µg/µl using Ultrapure water. RNA was transferred to 96-wells plate for easy downstream processing and stored at -80°C prior to use. The IVTsgRNA system as described here was validated and compared to Guide-it sgRNA In Vitro Transcription Kit (Takara Biosciences, 632635). Three different polymerases were validated to generate IVTsgRNA template: PrimeStar (Takara Bio Inc. R010A), Phusion High-Fidelity (Cat. #F553L,

Thermo Scientific™) or Taq polymerase (Cat. #10342046, Invitrogen). Nucleofection strips and cuvettes can be re-used by washing twice using 100 µl Ultra-pure water and by sterilized using 100 µl ethanol. Ethanol was removed from the strips after 1 minute of incubation and dried for 30 minutes in the flow hood.

**Poly chain reaction (PCR)**

For polymerase chain reactions (PCRs), 1-50 ng of DNA was amplified using Invitrogen Taq DNA Polymerase (Cat. #10342046) in 25 µl final reaction volume according to manufacturer's instructions. Samples were then amplified in Veriti 96-Well Thermal Cycler (Thermo Fischer), starting with 180 seconds of denaturation at 94°C followed by 40 cycles of denaturation (45 seconds, 94°C), annealing (30 seconds, 59-64°C) and extension (90 seconds per 1 kb, 72°C) and one final extension of 10 minutes at 72°C.

**PCR clean-up, sanger sequencing and indel analysis**

PCR cleanup was performed according to NEBs manufacturer's conditions by adding Exonuclease I (M0293) and Shrimp Alkaline Phosphatase (rSAP) M0371 to PCR products. Sanger sequencing was performed using the BigDye™ Terminator (BDT) Cycle Sequencing Kit (Thermo Fisher # 4337455) in a final reaction volume of 20 µl according to the manufacturer's instructions and sequenced in Applied Biosystems™ 3730 DNA Analyzer. Indel score were analyzed under standard conditions using TIDE analysis open software.[58]

### Supplemental figure legends

#### Supplemental Figure S1

**A**) General workflow Cas9 purification. 2-3 µg of Histrapp eluted fractions were ran on SDS-page gel (left panel), and Cas9 containing fractions were pooled and concentrated in 2-3 ml total volume. **B**) Size exclusion was performed on the pooled fractions to discriminate Cas9 monomers from oligomers to prevent protein aggregates in culture. Cas9 monomers typically elute around 66 ml (middle panel) and finally the purest fractions, indicated by SDS-page (right panel), were pooled and aliquoted snap frozen using liquid nitrogen and stored at -80ºC. (**C**) For concentration determination, we recommend to dissolve the BSA standard in at least 5x diluted SEC buffer, since NSDB-201, an important zwitter-ionic compound added to the pooled Cas9 fractions, interferes in the BCA assay (undiluted curve). (**D**) To check whether chaperone proteins are needed for active Cas9 expression, we compared parental BL21 cells (NEB) transformed with chaperone plasmids (Takara Biosciences Cat 3340), to Rosetta (DE3) cells. In both cases active Cas9 was purified, however, the yield of Rosetta cells is higher. (**E**) Next, we compared BL21-Star cells with Rosetta cells for protein production. Transformation efficiency of BL21-star cells was higher compared to Rosetta cells when 10 ng of DNA plasmid spCas9-3xNLS was used. (**F**) Expression of spCas9 from cultures induced at different OD's in LB media. In each lane 250 µl of cell culture was loaded, indicating that BL21-star cells provide higher yields compared to Rosetta cells and the optimal OD for protein induction is 0.6-0.7. (**G**) Data points from Figure 1A were plotted and correlated to the number of weeks past production to study stability of the buffer.

#### Supplemental Figure S2.

$CD34^+$ cells from 6 donors were nucleofected with indicated buffers and a 2xRNP targeting CD45**,** each dot represents a different donor and cell survival was measured at day 1 post nucleofection, and KO after 4 days. Statistics were calculated using a One-way ANOVA with Tukey's post hoc test (*<p0.05 ,**p<0.01, ***p<0.001, ****p<0.0001).

#### Supplemental Figure S3.

**A**) Illustration showing an overview of the PCR product that was generated to express sgRNAs. Gel images show the results from different primer concentrations and Taq-polymerases used. **B**) sgRNA yield after 16 hours of in vitro transcription. Each dot represents a sgRNA and error bars the SD. **C**) Comparison performance of IVTsgRNA products derived from Takara Bio kit and from our institute (left panel). Six distinct IVT were nucleofected in K562 and analyzed for Indels at indicated time points after nucleofection using TIDE analysis (right panel).

**D**) Validated sgRNAs from HSPCs were compared to the bulk of the in K562 tested IVTsgRNAs for 10 in silico prediction models present on CRISPOR website. Each data point represents a unique sgRNA. Student's t-test was used for statistical analysis (*<p0.05 ,**p<0.01, ***p<0.001, ****p<0.0001). **E**) validated sgRNAs from Figure 3B and Figure 3E are plotted for the Indel scores in HSPCs relative to their Wang *in silico* scores. P-value was calculated using Linear regression analysis.

**Supplementary Figure S4.**

Flow cytomertric analysis of gene edited HSPCs. Column total events P5 shows the number of events that were measured and analyzed in Figure 3B and Figure 3C. $CD34^{+}CD38^{-}CD45RA^{-}$ $CD90^{+}$ events, also referred to as the most potent $CD34^{+}$ HSCs.

**Supplementary Figure S5A-E. Cell line transductions and nucleofection optimization.**

**A)** W-blot enhanced image from figure 5B showing that ML2 cells are also weakly positive for p21 induction. **B**) Cell lines after transduction and FACS sorting, Dark grey shows that all cells, except NB4 were <90% for the BFP-GFP reporter. **C**) Cell line was nucleofected on different programs with 1 µg pMax-GFP plasmid. Bars represent cell viability and GFP positivity measured 24 hours post nucleofection. **D,E**) GFP reporters cell lines were nucleofected with 4 µg Cas9-3xNLS. Cell survival was measured 24 hours post nucleofection, knockout after 96 hours. Overall, these data suggest that X-01 can be used best HL60, whereas ML2 and EOL-1 perform better at CA-137.

**Supplementary Figure S5F-I supportive data for P21 induction experiments**

**F)** Cell survival $CD34^{+}$ cells measured using flow cytometry at indicated time points, data was analyzed using a Student's t-test. **G**) FACS plots belonging to figure 5G to show the KO efficiencies for each donor and indicated condition. **H,I**) showing the KO efficiencies for CD33 and CD45 in single and double KO. Note that CD45 sgRNA efficacy is significantly hampered in the double KO compared to single KO shown by Student's t-test analysis (*<p0.05 ,**p<0.01, ***p<0.001, ****p<0.0001).
