## supplemental figures for "Optimized guide RNA selection recommendations for using *sp*Cas9 gene editing in human hematopoietic stem and progenitor cells"

### Supplemental figure 1

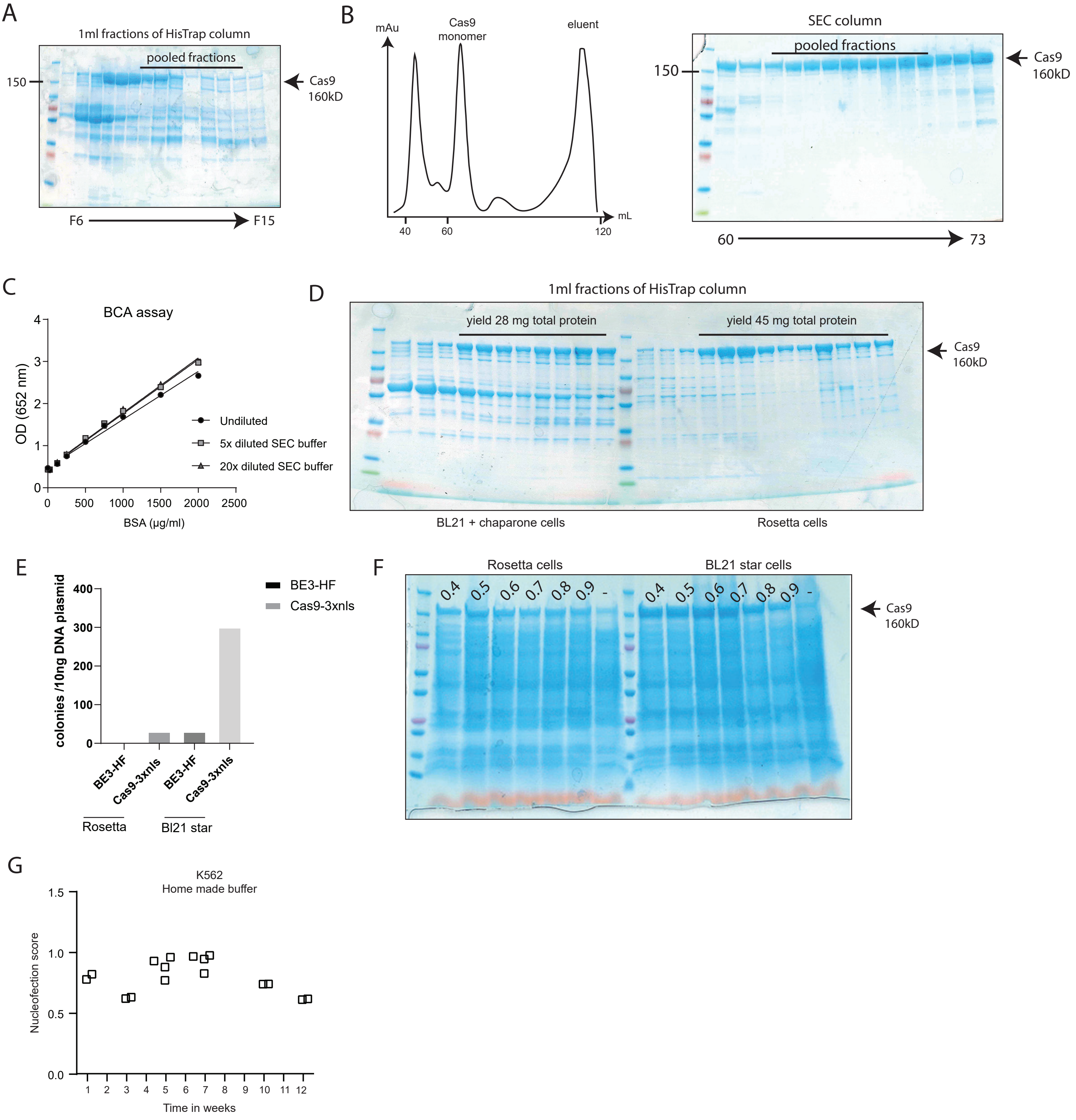

### Supplemental figure 2

A

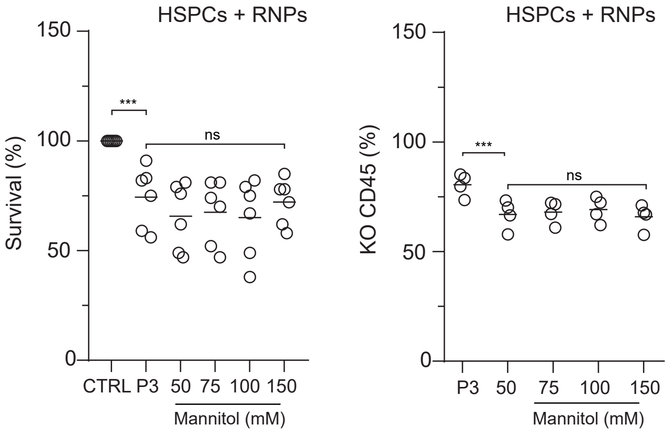

### Supplemental figure 3

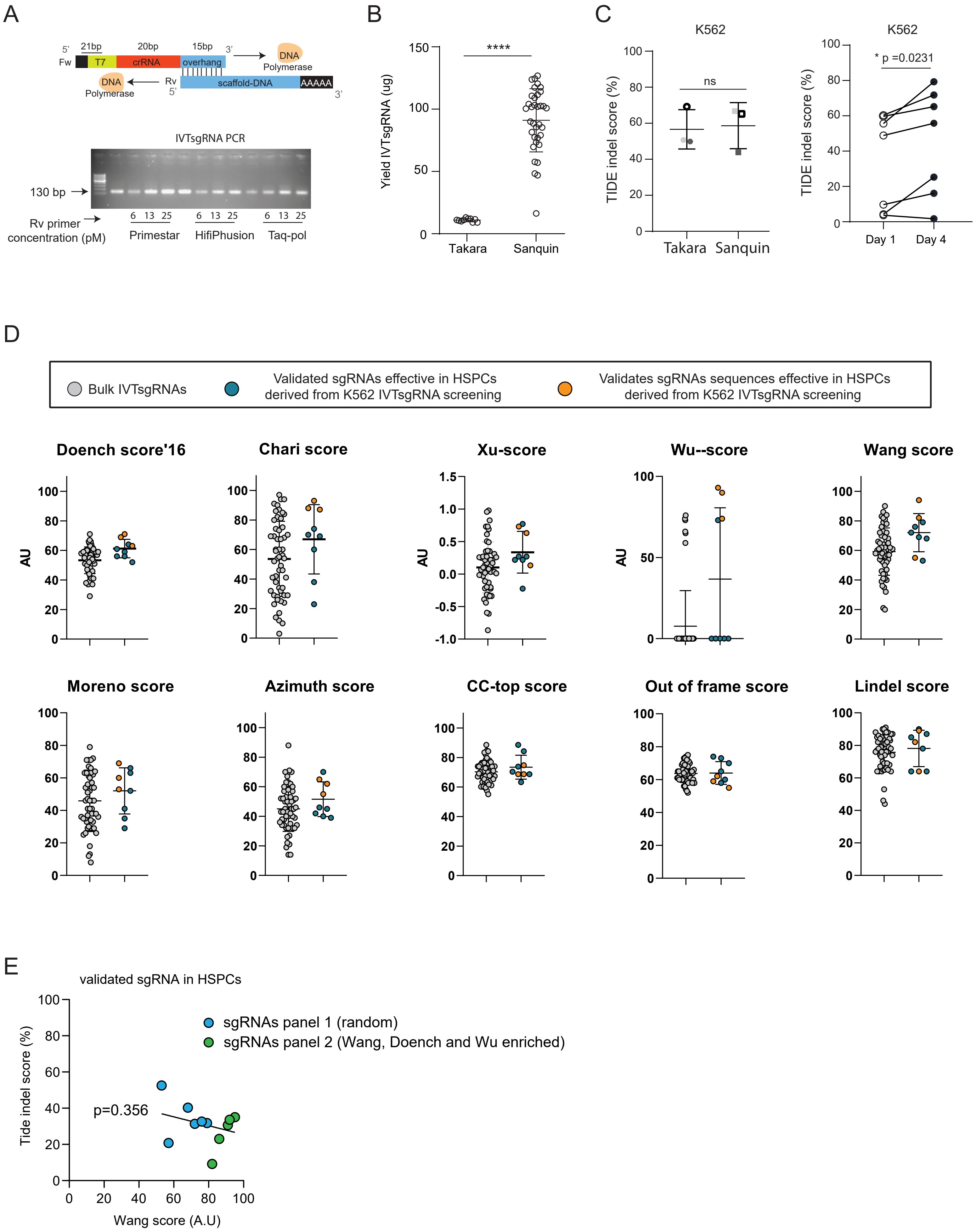

### Supplemental Figure 4

| CD34 Donor | conditions | total events measured | Total events P5 | % CD34+CD38-CD90+CD45RA- |
|---|---|---|---|---|
| | hCD33 | 15747 | 165 | 0,01 |
| | hCD45 | 13686 | 358 | 0,03 |
| | Cas9 Only | 35127 | 804 | 0,02 |
| 1 | No Nucleo | 29610 | 775 | 0,03 |
| | hCD33 | 16574 | 164 | 0,01 |
| | hCD45 | 19011 | 349 | 0,02 |
| 2 | Cas9 Only | 30305 | 861 | 0,03 |
| | hCD33 | 23784 | 1735 | 0,07 |
| | hCD45 | 28188 | 2885 | 0,10 |
| | Cas9 Only | 40493 | 4125 | 0,10 |
| 3 | No Nucleo | 40005 | 3438 | 0,09 |
| | hCD33 | 31725 | 1089 | 0,03 |
| | hCD45 | 28149 | 1257 | 0,04 |
| | Cas9 Only | 42370 | 2039 | 0,05 |
| 4 | No Nucleo | 38728 | 2240 | 0,06 |

### Supplemental figure 5

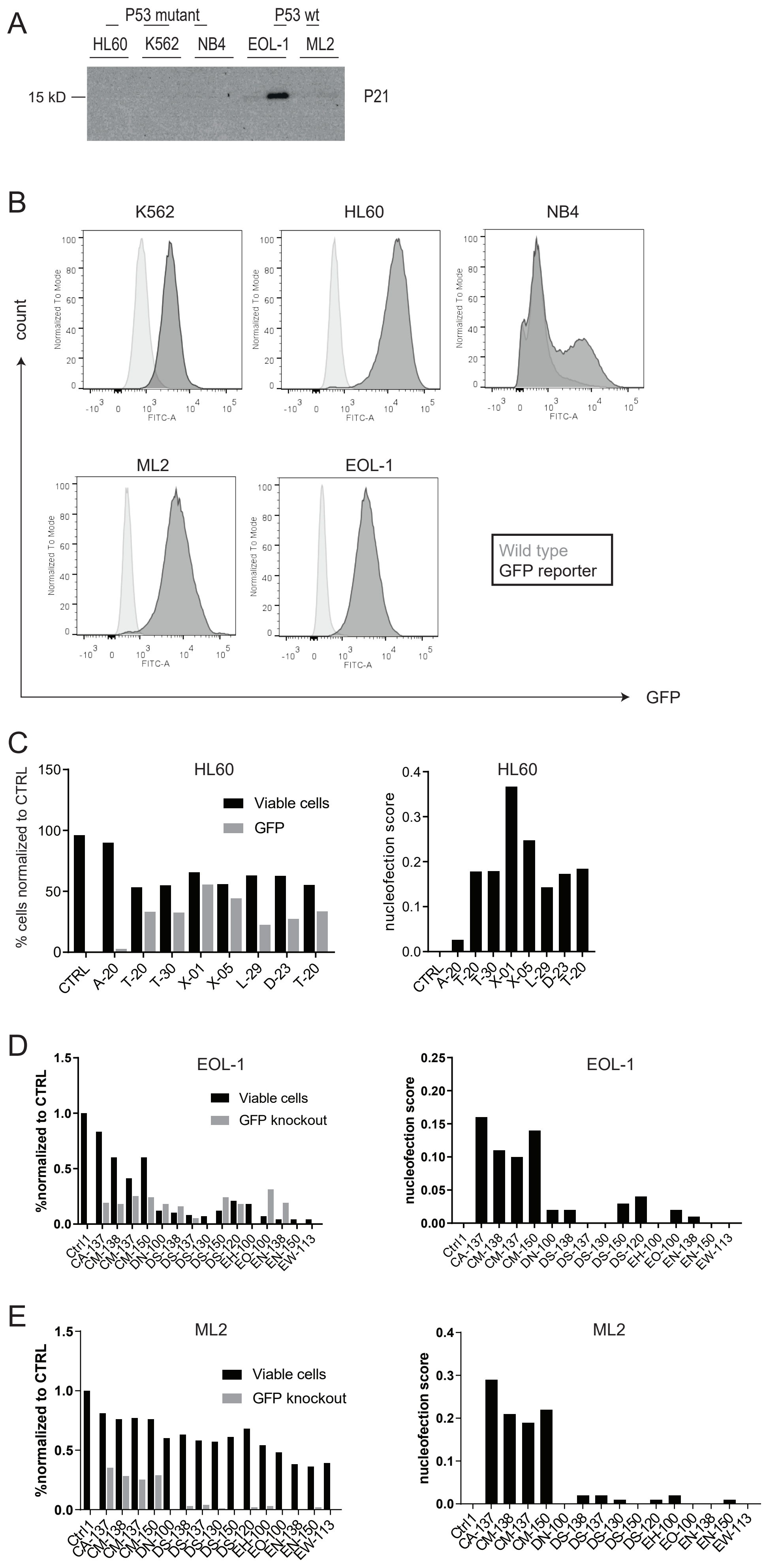

Supplemental Figure S5

F

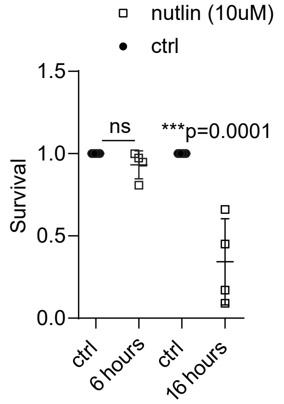

G

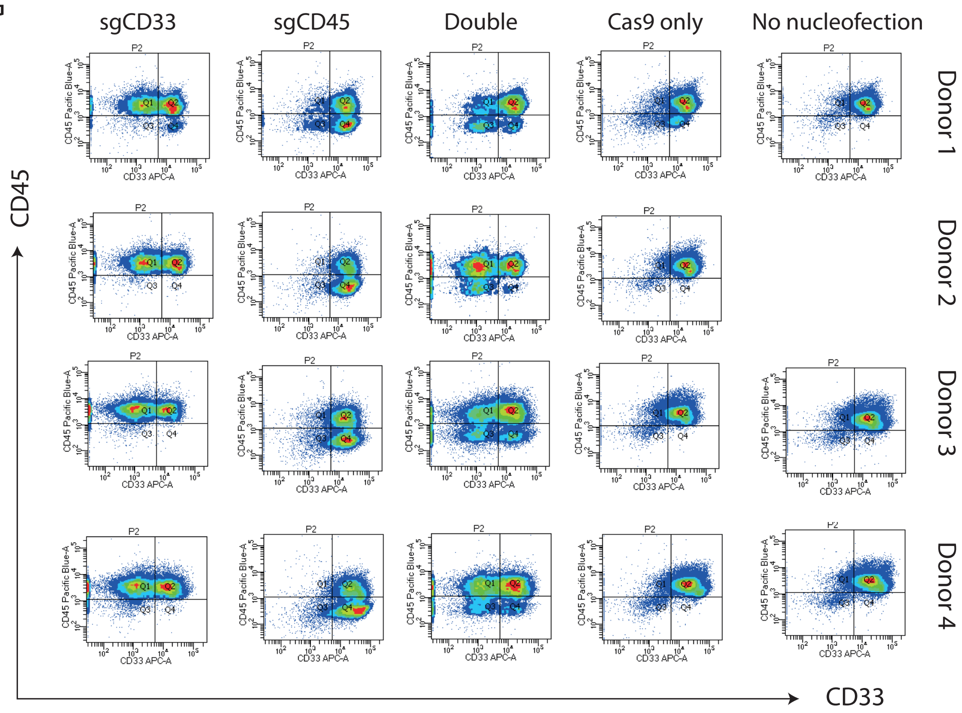

H

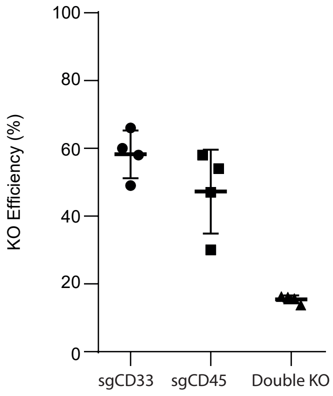

I

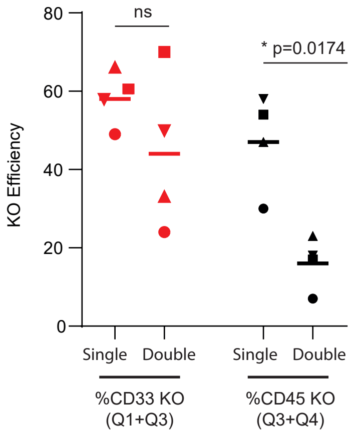
